# GPCRome-wide structural analysis of G-protein-coupling selectivity

**DOI:** 10.1101/2022.09.24.508774

**Authors:** Marin Matic, Pasquale Miglionico, Asuka Inoue, Francesco Raimondi

**Affiliations:** Laboratorio di Biologia Bio@SNS, Scuola Normale Superiore; Graduate School of Pharmaceutical Sciences, Tohoku University, Sendai, Miyagi 980-8578, Japan

## Abstract

We present a comprehensive computational analysis of available 3D GPCR-G-protein complexes to inspect the structural determinants of G-protein-coupling selectivity.

Analysis of the residue contacts at interaction interfaces has revealed a network of secondary structure elements recapitulating known structural hallmarks determining G-protein-coupling specificity, including TM5, TM6 and ICLs. We coded interface contacts into generic-number fingerprints to reveal specific coupling-determinant positions. Clustering of G_s_ vs G_i_ complexes is best achieved when considering both GPCR and G-protein contacting residues rather than separated representations of the interaction partners, suggesting that coupling specificity emerges as contextual residue interactions at the interface. Interestingly, G_s_-GPCR complexes contain a higher number of contacts than G_i/o_-GPCR complexes, likely caused by overall higher conservation and structural constraint on the G_s_ interface. In contrast, G_i/o_ proteins adopt a wider number of alternative docking poses on cognate receptors, as assessed via structural alignments of representative 3D complexes.

Furthermore, binding energy calculations demonstrate that distinct structural properties of the complexes contribute to higher stability of G_s_ than G_i/o_ complexes. AlphaFold2 predictions of experimental binary complexes confirmed several of these structural features and allowed us to augment the structural coverage of poorly characterized complexes (e.g. G_12/13_).

We propose that the structural properties of different G-protein complexes, such as structural restraining of G_s_ compared to G_i/o_ ones, could be instrumental in fine-tuning their activation and downstream signaling mechanisms.

**Highlights:**

-Comprehensive structural bioinformatics analysis of available GPCR-G-protein complexes captures common as well as group-specific structural features responsible of receptor-G-protein recognition
-Distinct contact patterns explain different docking modes of G_i/o_ vs G_s_ complexes, the latter being characterized by higher enrichment of characteristic contacts and lower structural variability suggestive of higher interface conservation.
-Structural hallmarks are associated with different estimated binding energies, which mainly discriminates G_s_ versus G_i/o_ couplings, but which also point to class-dependent differences (e.g. Class A vs Class B) in binding the same transducer (G_s_)

## Introduction

G-protein-coupled receptors (GPCRs) constitute the largest family of cell-surface receptors, making them a primary pharmacological class which is targeted by approximately one-third of the marketed drugs^1^. They transduce extracellular physico-chemical stimuli to intracellular signaling pathways by coupling to one or more heterotrimeric G-proteins^2,3^, which are grouped into four major families: G_s_, G_i/o_, G_q/11_ and G_12/13_ based on homology of their α-subunits^4^. GPCRs’ downstream activity is controlled by β-arrestins, which desensitize GPCRs’ activity and provide an additional layer of signaling modulation via ERK^5^. Ligand binding to GPCRs induces conformational changes that lead to binding and activation of intracellular G-proteins. Mammalian GPCRs display a wide and distinct repertoire of G-protein-coupling, ranging from highly selective to promiscuous profiles, which orchestrates specific downstream cellular responses^6^. Aberrant transduction mechanisms are linked to a myriad of pathological states (i.e. signalopathies), including cancer^7–12^. A deeper understanding of these signaling mechanisms, integrated in the wider biological context defining a disease state, can impact targeted therapies and personalized medicine protocols (e.g.^13^).

Determining specific coupling profiles is critical to understanding GPCR biology and pharmacology. On one hand, quantitative screening methodologies have been set up to systematically profile the binding activities of GPCRs for transducer proteins^14–17^. Based on these large scale experimental assays, sequence-based machine learning for coupling specificity have been proposed^18,19^.

On the other hand, the determination of receptor-G-protein complex structures is progressing rapidly, with over 260 complex structures deposited in the PDB (as of July 2022). The determination of the first structures of G_i_ coupled receptor complexes allowed for initial comparisons with G_s_ counterparts and highlighted a role of TM5 and TM6 as selectivity filter^20–24^. As a complement to the release of the MT1-G_i_ complex, we also systematically compared available G_s_ and G_i/o_ complexes with Class A receptors in terms of interface contact networks and G-protein docking mode similarity assessed via structural alignment^25^. The recent determination of four structures of the serotonin receptors (e.g. 5-HT4, 5-HT6 and 5-HT7 with G_s_, and 5-HT4 with G_i1_) confirmed the role of TM5 and TM6, and in particular their variable length, as a selectivity filter for G-protein binding^26^. The authors also showed via bioinformatics analysis that this macro-switch is conserved among other class A GPCRs^26^. Yet, a comprehensive pictures of the structural hallmarks of coupling specificity remains elusive.

In this study, we analysed through structural bioinformatics available GPCR-G-protein 3D complexes to shed further light on the structural basis of coupling specificity through analysis of interaction interface contact networks, G-protein docking modes and binding energies (Fig. 1A).

**Figure 1:**
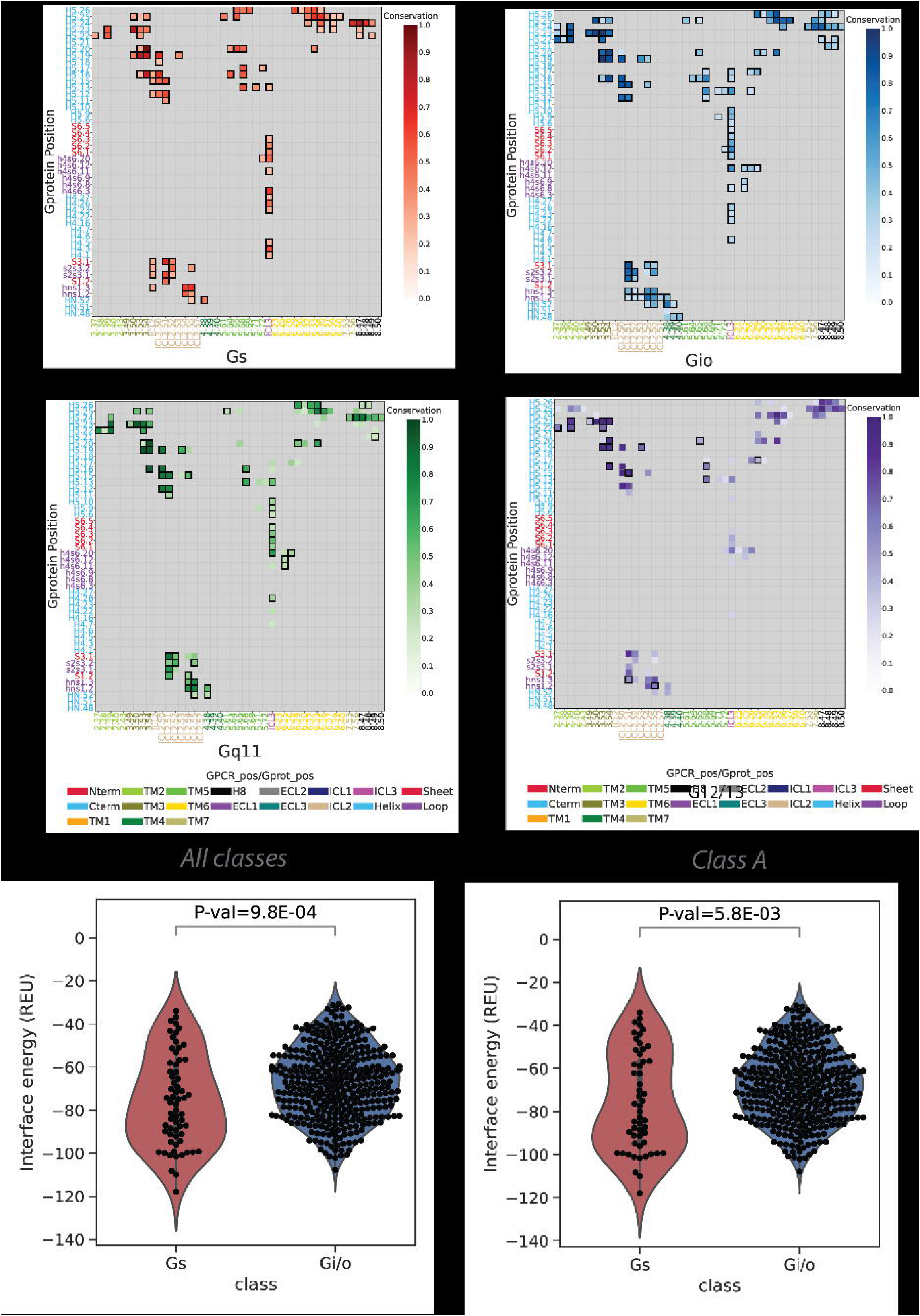
A) statistics of the total number of GPCR-G-protein complexes considered; B) number of representative GPCR-G-protein used for downstream analysis; C) workflow of the interface contact analysis procedure

## Results

### Different G-protein complexes are characterized by different contact network topologies

We considered a total of 264 3D experimental GPCR-G-protein complexes, comprising 126 G_s_, 133 G_i/o_, 4 G_q/11_ and 1 G12/13 complexes, corresponding to 63, 14, 1, 3 and 2 unique receptors from Class A, B1, B2, C and Frizzled, respectively, and entailing 8 different G-proteins (i.e. *GNAS, GNAI1, GNAI2, GNAI3, GNAT1, GNAO1, GNAQ, GNA13*) (Fig. 1B; Table S1). To avoid any bias due to redundant structures determined for the same GPCR-G-protein complex, we derived a set of 94 non-redundant 3D complexes by considering representative structures for each receptor-G-protein pair, using resolution and canonical sequence coverage as selection criteria (Figure 1C; see Methods). We first identified the residues that are in spatial contact at the GPCR - G-protein interaction interface (see Methods). We then mapped contacting residues to consensus numbering through GPCRdb^27^ (Table S2) and the common G-protein numbering (CGN)^28^ schemes (Table S3), respectively for GPCRs and G-proteins, and then we aggregated contacts on the basis secondary structure elements(SSEs), to yield a network of interacting SSE elements at GPCR-G-protein interfaces (Fig.1A). For the most abundant coupling groups (i.e. G_s_ and G_i/o_), we derived coupling group specific SSE contact networks by pooling contacts on the basis of the bound G-protein. SSE contact networks highlight structural signatures specific to each coupling group. Certain SSEs are invariably central within the interface network, such as TM5, ICL2 or ICL3 for GPCRs (Figure 2A-C) or H5 for the G-protein (Figure2A,B,D). Other elements vary their connectivity based on the bound G-protein. In particular, TM5 has a higher degree of contacting SSEs in G_s_ complexes as well as overall number of contacts, while G_i/o_ complexes are instead characterized by higher interconnectivity at the ICL1, TM6, ICL3 and H8 (Figure 2A,C). Differences in the overall network topology also emerged when we measured the information flow, quantified as the number of shortest paths passing through each node (i.e. betweenness centrality; see Methods). Indeed, TM5 and ICL2 have a higher betweenness centrality in G_s_ complexes, while TM3, TM6 and ICL1 prevail in G_i/o_ ones (Figure 2E). Overall, G_s_ contact graphs are significantly different from G_i/o_ ones, as assessed by comparing distances, computed as the Frobenius norm of the difference between the adjacency matrices of the interface contact graphs (permanova p-value= 1E-06; see Methods).

**Figure 2:**
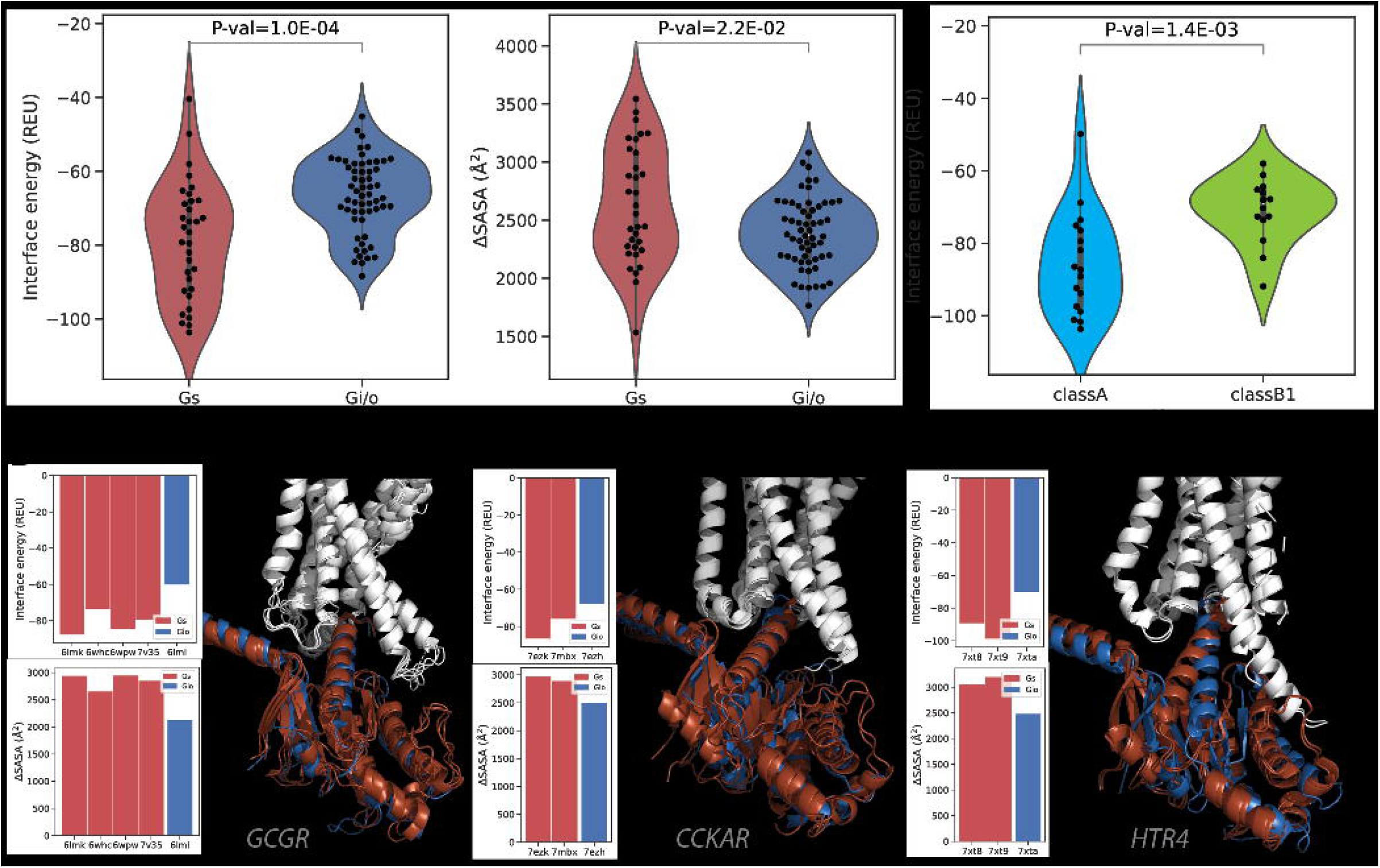
interface contact network analysis: A) SSE contact network for Gs1 complexes: GPCR and G-protein nodes are colored in green and cyan, respectively. Node diameter is proportional to the total number of contacts mediated by that SSE. Edge thickness is proportional to the number of contacts between connected SSEs and coloring (darker red) is directly proportional to contact conservation; B) SSE contact network for Gi/o complexes. Network characteristics as in 2A; C) GPCR SSE network node degree distribution for Gs1 and Gi/o networks; D) G-protein SSE network node degree distribution for Gs1 and Gi/o networks; E) GPCR SSE network betweenness centrality distribution

### Contact interface fingerprints imprint coupling specificity

We employed interface contacts to build interaction fingerprints, which are vectors that numerically encode the presence or absence of a contact and which can be used to compare in an unsupervised way GPCR-G-protein complexes based on their interface’s structural features (Fig.1A). We have generated interface fingerprints by mapping either residue pairs at each vector position (Complex fingerprints, or CF), or contact positions separately for the receptor and G-protein (Receptor and G-protein fingerprints, respectively RF and GF; see Methods). We also estimated the contact positions that are more frequent than expected in a given coupling group through log-odds ratio statistics (see Methods), and we used this information to filter the most informative contacts for G_s_ and G_i/o_ couplings (Figure 3A). CF clustering identifies two main clusters: the largest one (cluster 1), is enriched with G_i/o_ complexes and mainly involve class A receptors, though a minority of class B1, C and F representatives is also observed, forming a sub-cluster together with a few sporadic G_s_ complexes (Fig. 3A). The second cluster (i.e. cluster 2) is enriched with G_s_ complexes from both class A and Class B. In cluster 1, more receptors show promiscuous couplings towards other G-proteins, in particular towards G_12/13_ (31% vs ~7% in Cluster 2) and G_q/11_ (49% vs 39% in Cluster 2). The higher promiscuity between G_i/o_ and G_12/13_ couplings is also observed when considering all couplings from the Universal Coupling Map (UCM^29^; Fig. S1). Indeed, when relaxing the criteria to perform CF clustering, by removing the log-odds ratio filter and by including all unique complexes, we observed that available G_12/13_ and G_q/11_ complexes cluster within the largest group, which is enriched in G_i/o_ complexes and in secondary couplings for G_12/13_ and G_q/11_ (Fig. S2). In details, the only G_12/13_ complex structure available (i.e. *S1PR2-GNA13*, PDBID: 7T6B), is clustered together with the G_i/o_ complexes of G_12/13_ binders, such as *LPAR1 and S1PR5*, suggesting a structural (likely evolutionary imprinted - see Discussion) connection between G_i/o_, G_q/11_ and G_12/13_ proteins.

**Figure 3:**
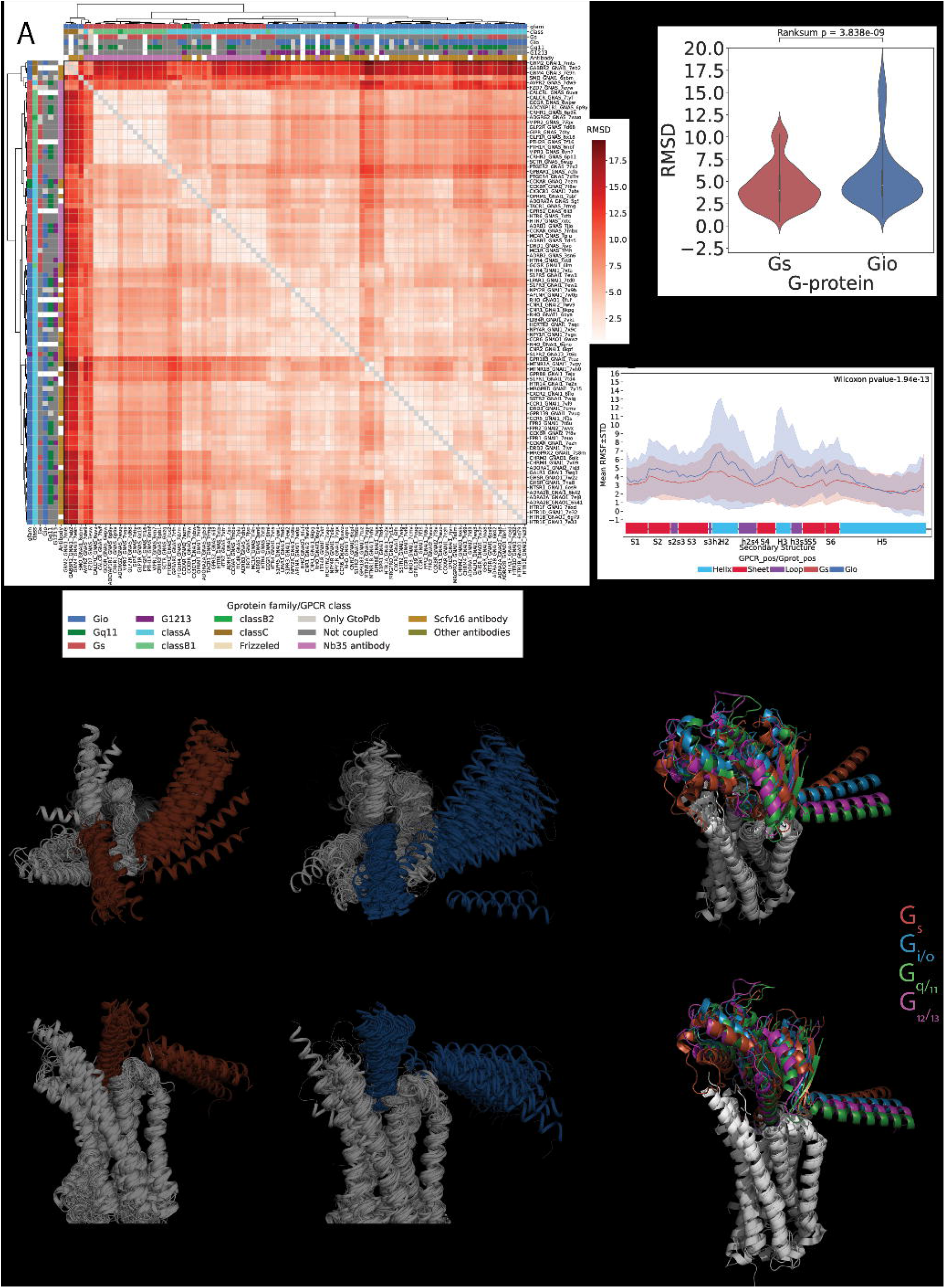
A) GPCR-G-protein contact interface fingerprint (or CF fingerprint): each row is a GPCR-G-protein contact positions (referenced respectively to GPCRdb numbering and G-protein position (CGN) numbering) and each column is a unique receptor complex. If a receptor is complexed with more than one G-protein, its complex fingerprint is reported accordingly. Columns are color annotated to indicate: G-protein bound in the experimental structure, GPCR class, experimental reported coupling (according to UCM, or either GEMTA, Shedding or GtoPDB). Right-side plot indicates the log-odds ratios (LORs) of the contacts observed at each position. Only contacts present in at least 10% of the structures and having an absolute LOR value greater than 2 are considered; B) fractions of experimental coupling groups of the receptors clustered identified through CF clustering; C) distribution of the LOR statistics for GPCR-G-protein contacts; D) Gi/o log-odds ratio statistics represented with a color scale ranging from blue (negative LOR) to red (positive LOR) using B-factor annotations on a representative structure (PDB ID: 7vl9): receptor (top, chain R), G-protein (bottom, chain A) along with distribution of the LOR statistics for GPCR (top) and G-protein (bottom) contact positions; E) Gs complexes contact frequency heatmaps: columns are GPCR positions in GPCRdb numbers, rows are G-protein positions in CGN numbers. Only contacts with frequency > 20% (over the number of unique complexes) are considered; F) structural comparison of Gs complexes mediated by a class A (*HTR6;* cyan; PDB: 7xtb) and classB representative (*ADCYAP1R1;* light green; 6p9y) and zoomed view of the contacts mediated by GPCR positions 6.39, 6.40 and 5.69, respectively with G-protein positions H5.23, H5.24, H5.25, H5.26; G) Gi/o complexes contact frequency heatmaps: representation features are as in 3E; zoomed view of the contacts mediated by GPCR position 3.49 in a representative Gi/o complex (*CCKAR*; PDB 7ezh) and distances between closest G-protein amino acids to E138 3.49 in the Gs complex of the same receptor (PDB: 7mbx).

Overall, G_s_ couplings are characterized by a significantly higher number of enriched contacts with respect to G_i/o_ ones (Pmann-whitney=1.19E-4; Fig. 3C). We also performed clustering and enrichment with fingerprints of receptors (RF) and G-proteins (GF) separately. The RF clustering chiefly points to inter-class differences, separating complexes formed by ClassA receptors from those involving other classes (Fig. S3A) while not showing particular contact enrichment differences between G_s_ and G_i/o_ complexes (Pmann-whitney=3.5E-1; Fig.3D). The GF clustering better separates these groups (Fig. S3B) and displays greater differences in contact enrichment distributions, though not yet significant (Pmann-whitney=3.5E-1; Fig. 3D). This suggests that the combination between G-protein and receptor’s residues provides maximum fine tuning to the recognition process (Fig. 3C).

Complex fingerprints clustering and contact heatmaps helped visualizing the contact positions that are characteristic of certain G-protein-couplings (Fig.3A, E, G). For instance, the following couplings are exclusively enriched in G_s_ complexes: ICL2.52-G.S3.1, 5.64-G.H5.20, 5.76-G.h4s6.3, 6.32-G.H5.24, 6.39-G.H5.24, 6.40-G.H5.25. The latter positions (i.e. 6.39 and 6.40) are example of G_s_, class B specific contacts (Figure 3A and Fig. 3A). These are favored by the characteristic TM6 break characterizing Class B receptors^30^, which allows residues on the TM6’s N-terminal half to approach the G-protein H5 C-term. It must be noted however that a few class A receptors also display contacts mediated by position 6.40, either with G_s_ (*DRD1* and *AVPR2*) or G_i/o_ (*MTNR1A* and *ADRA2A*). Other G_s_ contacts, such as 5.76-G.h4s6.3 and 6.32-G.H5.24 appear to be instead class A-specific (Figure 3A). On the other hand, the following contacts are exclusively enriched in G_i/o_ complexes: 2.39-G.H5.24, 3.49-G.H5.23, 3.50-G.H5.24(or G.H5.25), 3.53-G.H5.20 (or G.H5.22, G.H5.23). Particularly striking is the enrichment of the contact involving the highly conserved D^3.49^, which is found exclusively in G_i/o_ complexes (Fig. 3A, G, H), while the DRY R^3.50^ or 3.53 positions also participate to universal contacts characterizing both G_i/o_ and G_s_ complexes (e.g. 3.50-G.H5.23, 3.53-G.H5.16, 3.53-G.H5.19; Fig. 3E,G). Other contacts specifically enriched in G_i/o_ complexes are: ICL2.51-G.hns1.3 (ICL2.51-G.s2s2.3) ICL2.54-G.S3.1, 5.71-G.H5.9, 6.25-G.h4s6.9, 6.28-G.h4s6.9, 6.29-G.H5.17, 6.29-G.H5.26, 6.33-G.H5.20 (G.H5.25) as well as all the contacts mediated by positions 8.47, 8.48, 8.49, 8.50 (Fig.3A,G).

Overall, GPCR positions such as 3.55, 5.64, 5.65, 6.68, 5.69, 5.72, 5.74, 5.76, 6.39 and 6.40 are enriched in G_s_ complexes(Figure 3D,E), while position 2.37, 2.39, 3.49, ICL2.50-2.55, 6.25, 6.26, 6.29, 6.33 or 8.50 are enriched in G_i/o_ complexes (Figure 3D,G). Likewise, the G-protein contact positions specifically enriched in G_s_ complexes are h4s6.20, h4s6.3, H4.3, H4.26, H5.11 (Figure 3D,E), while contact positions h4s6.9, h4s6.12, s6.1, H5.9, H5.21, H5.22, H5.26 are enriched in G_i/o_ complexes (Figure 3D,G).

Notably, certain GPCR positions hold switch characteristics, in other words some of the contacts that they mediate are enriched in G_s_ and others in G_i/o_ depending on the partner residues. For example, the contact of 5.65 with G.H5.16 is enriched in G_i/o_, while the ones with G.H5.25 and G.H5.26 in G_s_. Similar patterns are observed for distinct contacts mediated by positions 5.68,5.69 and 5.72 (Fig. 3A, E,G).

### Different repertoires of G-proteins docking modes

We assessed the overall structural similarity of GPCR-G-protein complexes via structural alignment, with a particular focus on the docking mode similarity of the G-protein α-subunits with respect to the receptor. To this end, we first superimposed the Cα-atoms of the most conserved positions within the 7TM bundle (i.e. that are present in all the solved structures), and then calculated the Root Mean Squared Deviation (RMSD) of the Cα-atoms of conserved positions of the Gα subunit (Fig.1A; see Methods). The clustering of 3D complexes based on their RMSD shows that G_s_ complexes tend to cluster separately from G_i/o_ ones (Pmann-whitney=3.8E-9; P-permanova=1E-6; Figure 4A,B and Fig. S4), being also different in variance with G_s_ complexes having a smaller variance (P-permdisp=0.034). When considering only class A receptors, the RMSD distributions are no longer significantly different (Pmann-whitney=5.5E-1; Fig. S5), although characterized by significantly different centroids (P-permanova=1E-6). RMSD clustering recapitulates certain structural hallmarks already emerged through interface analysis. Indeed, the largest cluster comprises only Class A receptors, the vast majority bound to G_i/o_ proteins, with the only exception of a subcluster comprising few G_s_ complexes (i.e. *ADRB1,2,3, DRD1, CCKAR, MC1R, MC4R, GPR52, HTR4, HTR6, HTR7*) and the *S1PR2-GNA13* complex. The second largest cluster involves almost exclusively G_s_ complexes, including a few Class A and the totality of class B receptors. This cluster also comprises two G_q_ complexes (i.e. *CCKAR* and *CCKBR*) and two G_i/o_ complexes (i.e. *OPRM1* and *CX3CR1*). Finally, a third, smaller outgroup cluster contains exclusively G_i/o_ complexes involving class A, C and F receptors most deviating from the other structures. Also in this case, the receptors in the largest cluster show a more promiscuous tendency, with G_q/11_ and G_12/13_ as most recurrent secondary couplings and G_s_ as the least recurrent one (Figure 4A). We have also estimated residue level deviations of the Galpha subunits of the fitted complexes by calculating Root Mean Square Fluctuations (RMSF; see Methods) and compared the profiles obtained for G_s_ and G_i/o_ complexes, which highlighted overall higher fluctuations for G_i/o_ complexes with respect to G_s_ ones, for the whole RasGTPase domain and in particular for regions such as H2 and H3 (Fig. 4C). Overall, G_s_ complexes display less variability of the terminals, with H5 appearing more conformationally restrained and bent towards TM3 and ICL2, while G_i/o_ complexes display greater conformational variability for αN and H5 (Fig.4D left). Comparison of representative structures of the four coupling groups show slight differences in the docking mode of each representative, which are nevertheless smaller for G_i/o_, G_q_ and G_12/13_ (Fig.4E).

**Figure 4:**
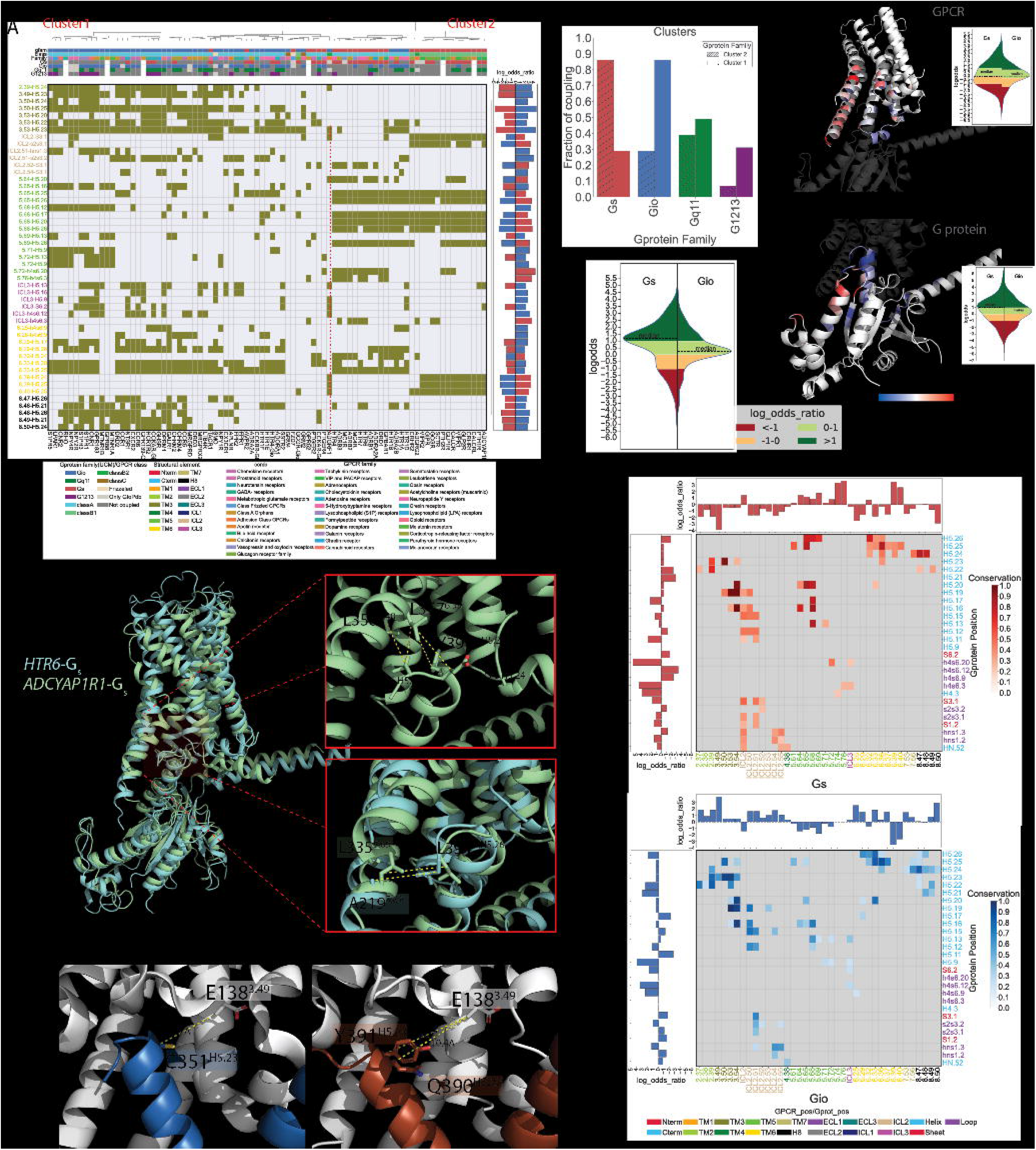
measuring similarity of G-protein docking modes: A) RMSD hierarchical clustering. Color annotations are as in Figure 3B with additional last field containing the information regarding presence of nanobodies; B)distribution of the RMSD within Gs and Gi/o complexes; C) root mean squared fluctuations of the G-protein consensus positions; D) superposition of class Gs and Gi/o representative complexes: GPCR 7TM bundles are represented as white cartoons; the N-term and C-term of the Gs (red, left) and Gi(blue, right) alpha subunits are represented as marker of the G-protein structural variability on experimental complexes; E) structural superimposition of representative structures defined on the basis on minimum RMSD to other members of the group (for G_s_ and G_i/o_) and release date (G_q/11_): G_s_(PDB=7xtb; red), G_i/o_ (PDB=7vl9; blue), G_q/11_(PDB=7ezm;gree), G_12/13_(PDB=7t6b;purple).

We also explored the potential conformational bias of the nanobodies (i.e. Fab16 or Nb35) used to stabilize the bound G-protein on the observed G-protein docking modes. First, we annotated the presence/absence of the nanobody for each complex subjected to RMSD clustering. We observed no correlation between RMSD clusters and the presence or absence of nanobodies (Fig. 4A, Fig. S4). Second, we relaxed the GPCR-heterotrimeric G-protein complex without such nanobodies, using state of the art methods for structural refinement (Rosetta relax; see Methods). Both RMSD and RMSF analysis performed on relaxed structures showed even larger statistically significant differences between G_s_ and G_i/o_ complexes (Pmann-whitney=2.71E-29; Fig. S6). Notably, the differences of the G_s_ and G_i/o_ RMSD distributions is still significant when considering only class A receptors (Pmann-whitney=6.2E-8; Fig. S7).

### Different energies characterize specific GPCR-G-protein interfaces

We exploited the relaxed GPCR-heterotrimeric G-protein complexes to further characterize the binding interface energy of the complex using Rosetta InterfaceAnalyzer^31^ (see Methods). By considering all available GPCR Gα-subunit pairs with an experimentally resolved complex, we showed that the ΔG of binding of Gs complexes is significantly lower than G_i/o_ complexes (Pmann-whitney=1E-4; Fig. 5A) and it partially correlates with the slightly higher ΔSASA observed for G_s_ complexes compared to G_i/o_ ones (Pmann-whitney=2.2E-2; Fig. 5B). When considering class A receptors only, the difference in binding energy distribution between G_s_ and G_i/o_ complexes is even larger (Pmann-whitney=1.1E-5; Fig. S8). Intriguingly, we observed that G_s_ is bound less strongly in class B than class A receptors (Pmann-whitney=1E-3; Fig. 5C), suggesting that receptors from different classes might bind to the same G-protein with different affinities due to different structural and functional requirements. On the other hand, the same receptor always binds with higher affinity to G_s_ than G_i/o_. Indeed, we have compared the binding energies of complexes of the same receptor (e.g. *GCGR*, *CCKAR* and *HTR4*) with both G_s_ and G_i/o_ proteins. Notably, the ΔG of binding for G_s_ is always lower and is characterized by higher ΔSASA compared to G_i/o_ irrespective of the slight docking modes variations observed in G_s_ complex structures of the same receptor (Fig. 5 D-F).

**Figure 5:**
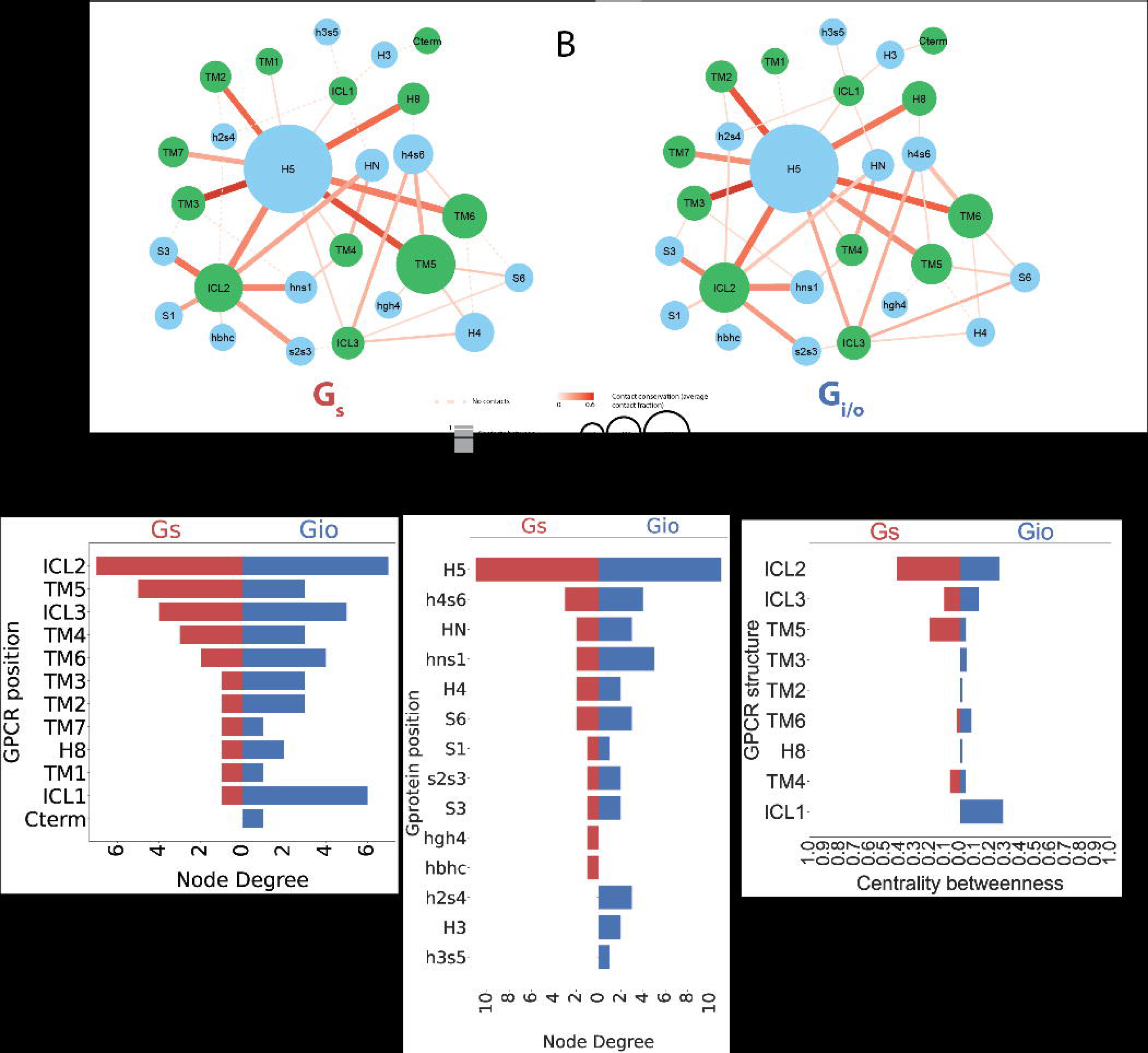
binding energy estimated through Rosetta A) ΔG binding (RU); B) Delta Solvent Accessbile Surface Area (ΔSASA); C) ΔG binding (RU) of Gs complexes with classA and classB receptors; Interface energy, ΔSASA and superimposed 3D cartoon models of Gs (red) and Gi/o (blue) complexes of D) *GCGR*; E) *CCKAR;* F) *HTR4*.

### AlphaFold2 predictions extend our understanding of the structural basis of coupling specificity

To further our understanding of the structural basis of GPCR-G-protein recognition, we predicted through AlphaFold-multimer^32^ 996 GPCR - G-protein alpha subunit pairs pairs from UCM binary interactions (see Methods). After filtering the predicted complexes based on structural, topological and energetic characteristics (see Methods), we ended up with 685 3D complexes for downstream processing (Table S4). Contact analysis performed on predicted complexes revealed patterns similar to those observed experimentally, in particular, the models recapitulated extremely well the experimental contacts of G_s_ and G_i/o_ complexes (Fig. 6A,B), likely due to abundance of experimental structures for these couplings. As expected, the agreement of contact patterns from experimental and predicted G_q/11_ and G_12/13_ complexes is lower due to fewer structural templates available for this coupling groups(Fig. 6C,D). Notably, contact heatmaps derived from each G-protein group display highly specific patterns, which could potentially illuminate novel signalling mechanisms for poorly studied G-proteins such as G_12/13_. For instance, contact frequency heatmaps for G_12/13_ complexes display peculiar patterns at the TM5 and ICL3, lacking contacts observed for other G-protein complexes such as 5.61-G.H5.25 and 5.63-G.H5.25, as well as several contacts btween ICL3 and G.H4 (Fig. 6D). Moreover, we also confirmed on predicted structures the differences in binding energies observed between G_s_ and G_i/o_ complexes, which are significant by considering complexes from either all classes (Pmann-whitney=9.8E-4; Fig.6E) or just class A (Pmann-whitney=5.8E-3; Fig. 6F).

**Figure 6:**
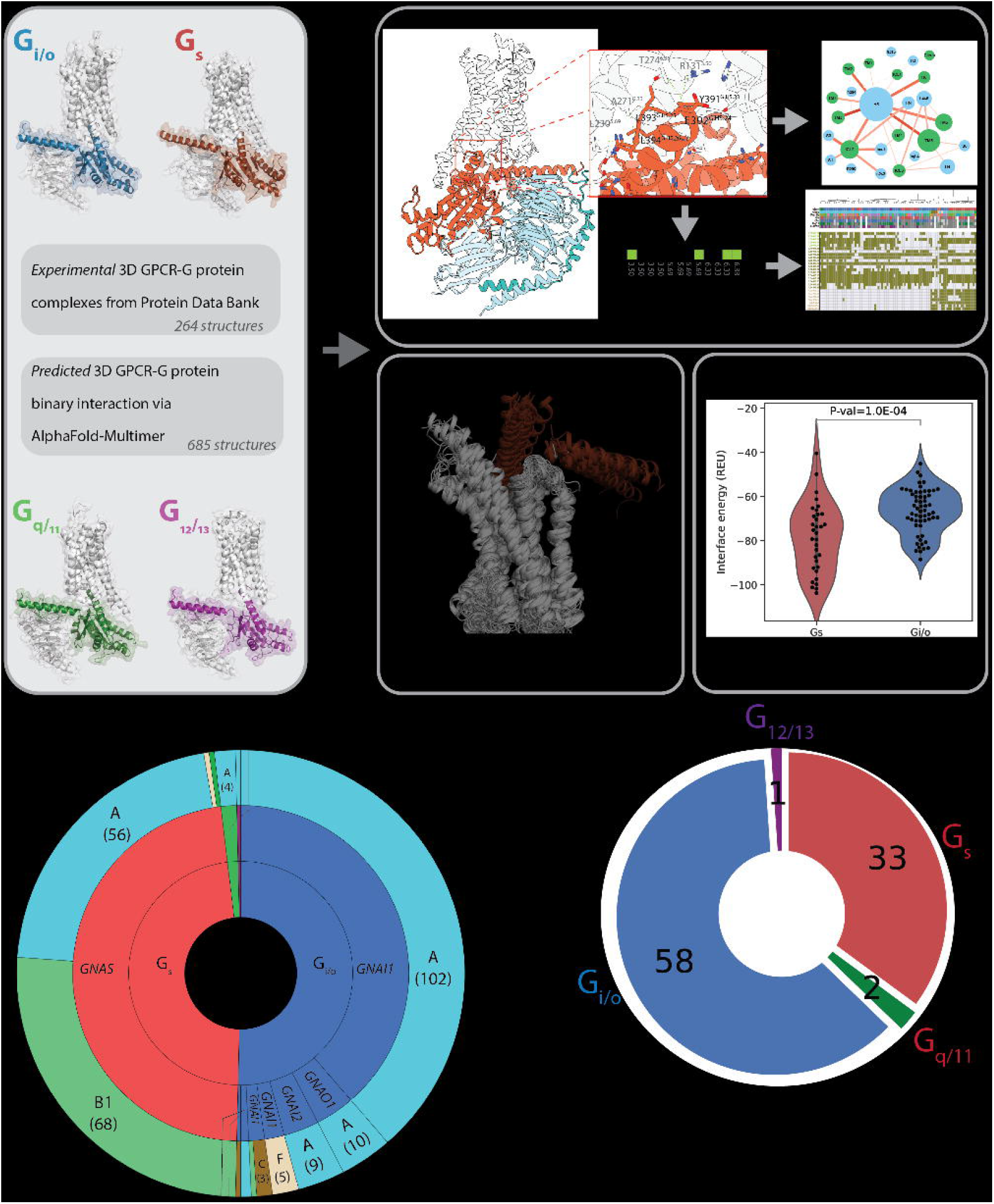
complexes contact frequency heatmaps of AlphaFold-multimer predicted complexes with A) G_s_, B) G_i/o_, C)G_q/11_, D)G_12/13_. Experimental contacts are marked with a dark square line; predicted interface binding energies for AlphaFold-multimer predicted complexes forE) all GPCR classes and F) only class A receptors.

## Discussion

In the present study, we have performed a computational, comparative analysis of 3D GPCR-G-protein complexes to identify structural hallmarks linked to coupling specificity.

Complexes involving different G-proteins are characterized by distinctive structural signatures, such as contact networks displaying a different engagement of secondary structural elements such as TM5, TM6 and ICLs. This notion is in line with the comparative analysis highlighting a selectivity filter operated by TM5 and TM6 (commented on^24^). More recently, structural determination of serotonin receptors in complex with either G_s_ or G_i_ proteins, accompanied by a bioinformatics analysis of a representative set of class A complexes, supported the role of these secondary structure element by highlighting a macro-switch, operated by TM5 and TM6 terminals’ variable length, which dictates selectivity towards G_s_ vs G_i/o_ ^26^. Our unsupervised analysis of interface contacts entails complementary interactions between key positions on both the receptor and G-protein sides. G_s_ complexes are characterized by a significantly higher fractions of enriched contacts, that are mainly imposed by contacting positions on the G-protein’s side. However, the combination between GPCR and G-protein residues leads to a maximum discrimination between the G_s_ and G_i/o_ groups. These results are reminiscent of the G-protein barcode model for coupling specificity, which emerges by the presentation of an evolutionary more rigid G-protein barcode to a more flexible receptor counterpart^33^. At the same time, G_s_ bound receptors are characterized by a less promiscuous binding, suggesting that the structural requirements for G_s_ specific binding overall impose a discrimination for other secondary couplings.

Certain SSEs are exclusively characterized by contact enrichments for specific G-protein (e.g. TM2,TM3 or H8 for G_i/o_). Notably, the DRY motif mediated contacts, particularly D^3.49^’s one, can be considered one of the main structural hallmarks of G_i/o_ vs G_s_ complexes. The latter are indeed characterized by a greater bending of G-protein H5 towards TM3 C-term and ICL2, which function as hinge to concomitantly detach the G-protein C-term from the DRY motif. Other regions are characterized by a switch-like character, meaning that certain positions form contacts enriched in G_i/o_ and others in G_s_, including: ICL2, which encompasses the known ICL2.51 specificity determinant position^34^, TM5, ICL3 and TM6. We noted that the switching behaviour might pertain also to individual positions and depend on the specific contact partner. This features are likely connected to the macro-switch described for the serotonin receptors ^26^.

Through our comprehensive analysis, we show that the few G_q/11_ and G_13_ complexes available display contact fingerprint characteristics similar to G_i/o_ complexes. These structural properties correlate with recent phylogenetic analysis showing that G_i_ and G_q_ family members share a common ancestor, and that G_12/13_’s ancestor is likely a retro-gene originated by retroposition from a pre-G_q_ gene^35^. The usage of state-of-art AI model (i.e. AlphaFold-multimer^32^) for structural prediction also allowed us to expand the structural repertoire of GPCR-G-protein complexes. This is particularly valuable for poorly characterized groups, such as G_12/13_ ones. Indeed, we predicted peculiar contact patterns at the TM5 and ICL3 that are characteristic of this group and might suggest unique structural requirements. Indeed, the critical importance of this regions also emerged in our previous effort to engineer a G_12_-DREADD, which was achieved by swapping shorter ICL3 loops from GPR183 or GPR132 on hM3D and experimentally validated to be functional^14^.

The observed structural differences are linked with different predicted binding affinities for the distinct coupling group complexes, with G_s_ complexes being more stable than G_i/o_ ones. Slight differences in interface contacts and docking mode might reflect the difference in binding affinities for G_s_ observed for class A vs. class B receptors. We speculate that the higher binding energy of class B receptor, suggestive of less stable binding to G_s_, might be related to slower rates for G-protein activation observed for a representative class B receptor (*GCGR*) compared to class A one (*ADRB2*)^30^. On the other hand, comparison of the binding energies of G_s_ and G_i/o_ complexes also revealed significant differences, which hold true even when considering the complexes with the same receptor, further stressing the major contribution from the G-protein residues in priming the binding. We speculate that G_s_ has a higher energetic barrier for activation due to its peculiar biological role of AC activation, which should be tightly regulated. The conformational restriction in G_s_ complexes might contribute to spatio-temporally fine tune AC activation. On the other hand, looser G_i/o_ binding to cognate receptors could explain their reported faster nucleotide turnover^36^. Moreover, lower structural conservation and less affine G_i/o_ complexes are likely connected to the success of this coupling^14,29^, which is instrumental in providing a redundant mechansim for AC inhibition.

To conclude, the greater availability of experimental complex structure and increasingly accurate predicted models will allow us to better understand the structural basis of G-protein-coupling specificity in the future. This knowledge will be key to design better biased drugs, able to modulate only certain transducers, as well as it will be leveraged to improve the design of novel chemogenetic probes such as DREADD.

## Methods

### Data sets

We used the mapping between PDB^37^ and Pfam^38^ provided by SIFTS ^39^ (2022/07/23 update) to retrieve all the structures containing both a GPCR and a G-protein alpha subunit. We used the Pfam entry PF00503 to identify structures of G-alpha subunits, and Pfam entries PF00001 (rhodopsin receptor family - class A), PF00002 (secretin receptor family - class B), PF00003(class C receptors), PF01534 (Frizzled/Smoothened family) to identify GPCRs. We found 276 structures that met this criterion. If more than one GPCR or Gα chain were in the structure, we considered the pair of chains with the highest number of contacts between them (Table S1).

We considered as contacts all the pairs of amino acids with a distance of less than 8 Å between their Cβ (C□ for glycin - see below), following standard practices employed for contact analysis in structural predictions^40^. Only the structures in which all the contacts between the GPCR and Gα chains were mapped to the same pair of Uniprot accessions (according to SIFTS xml residue level mappings) were kept for further analysis, the other structures were excluded, considering them as chimeric. In this way, we were left with 264 structures.

Whenever we found more than one structure representing the same GPCR-G-protein pair, we considered the structure with the highest resolution to be the representative for each pair.

In case more than one structure had the same resolution, we chose the structure which covered the largest portion of the corresponding G-protein according to SIFTS. We found 94 GPCR-Gprotein pairs with at least one solved structure (Table S1).

### GPCR-Gα complexes prediction via AlphaFold-Multimer

A total of 996 GPCR-G□ pairs, reported to bind in UCM^29^, were considered, respectively corresponding to 164 and 13 human GPCRs and Gα proteins (the three members of the GNAT family are not considered). We generated through Alphafold-Multimer v2.2^32^ the 3D structural models for each of these experimental GPCR-Gα complexes lacking a known 3D structure in the PDB. The databases required to run AlphaFold-Multimer were downloaded on 15 March 2022. Among the 5 models generated for each GPCR-Gα pair, only the one with the highest confidence was considered for further analysis.

We removed 114 predicted complexes (882 structures left) based on those structure having Nterm or Cterm positions of GPCR in contact with part of G-protein, structures not having any H5 position of Gprotein in contact with part of GPCR, and no contact of GPCR with specific conserved positions of H5 in Gprotein (H5.16,H5.19,H5.20,H5.23,H5.24,H5.25). To further remove low quality models produced by Alphafold, we used the output of Rosetta InterfaceAnalyzer^31^(see below paragraph “Analysis of GPCR-Galpha binding energy with Rosetta”) and we kept only models with a ΔG < 0 REU, to avoid unnatural complexes with a positive binding energy, and with 1500 Å ≤ ΔSASA ≤ 3700 Å, which is approximately the range in which all experimental structures fall. This led to a total of 685 filtered, higher quality complex structures for downstream analysis (Table S4).

Moreover, we employed Alphafold’s confidence score (predicted Local Distance Difference test - pLDDT), to remove protein terminals predicted with low confidence which might lead to artefactual contacts. Hence, we trimmed the N-terminal and the C-terminal of both the GPCR and the G-protein up to the first residue with pLDDT > 70. We then performed the interface analysis on these trimmed sequences using the same procedure used for experimental complexes.

### Contact analysis

We considered residue-residue contacts mediating GPCR-G-protein interfaces as those having the Cβ spatially closer than 8 Å (C□ for glycin) (as in ^40^). We analyzed 264 solved PDB GPCRs-Gprotein complexes, which according to G-protein family classifcation comprised 133 G_i/o_, 126 G_s_,4 G_q/11_ and 1 is G_12/13_ structures. We mapped interface PDB residues to Uniprot canonical sequences residues by using SIFTS residue level mappings from individual PDB xml files. The interface sequence positions were then mapped to GPCRdb numbering^27^ (Table S2) and to the Common Gprotein Numbering (CGN) schemes^28^ (Table S3). We aggregated contacts of different structures referred to the same GPCR-G-protein complex and considered equivalent residues pairs from different structures only once to avoid redundancy. We compared G_i/o_ and G_s_ complexes by creating a consensus list of GPCR and G-protein positions found in contact in at least one of the members of the two groups. For each GPCR-G-protein consensus positions, we calculated the fraction of GPCRs displaying a contact in each G-protein family group (either G_i/o_ or G_s_) over the total number of GPCRs in contact. Such a fraction reflects the conservation of a contact in a given G-protein group. We constructed an interface contact network by considering the consensus list of GPCR-G-protein contacts from all complexes, as well as from only those including class A GPCR. Contacting positions were projected to the secondary structure elements of both GPCRs and G-proteins, represented as nodes of the networks. Nodes diameter is proportional to the total number of contacts mediated by the position of that secondary structure element. Edge width is proportional to the number of unique GPCRs mediating contact between the two linked SSEs. Edge coloring (bright to dark red) is proportional to the average of the contact fraction of individual position pairs, and it reflects the overall contact conservation between two SSEs. Dashed lines indicate contact not formed in a given G-protein class but present in the other. We calculated network statistics such as node degree and centrality betweenness distribution through Cytoscape (https://cytoscape.org/)^41^ and customized Python scripts. All the analyses have been done through customized Python scripts, using biopython libraries (version 1.78) and are available upon request. Network drawings have been generated through Cytoscape.

To analyze the statistical significance of the difference between G_s_ and G_i/o_ contact fingerprints, we generated an inter-chain contact graph for each structure. We then used the Frobenius norm of the difference between the adjacency matrices of the interchain contact graphs as a distance metric for the structures. The resulting distance matrix was used to perform a PERMANOVA test^42^ to determine if the graphs generated by interaction with the G_s_ and the G_i/o_ family were significantly different from each other. This test compares the difference in docking modes in the same G-protein family to the difference in docking modes between different G-protein families. To evaluate the difference in variance between the distribution of intrafamily pairwise distances, we used the PERMDISP test. Both tests were performed using the scikit-bio library (version 0.5.4) in python {https://cran.r-project.org/web/packages/vegan/index.html}. We performed 10^6^ and 10^4^ permutations for PERMANOVA and PERMDISP, respectively.

Analysis was also done on Alphafold predicted complexes considering 362 G_i/o_, 212 G_q/11_, 64 G_s_ and 47 G_12/13_ structures according to UCM. Networks have been created for full range of position pairs and for cases where >0.2 complexes have the position pairs.

### Fingerprint analysis

We have generated interface fingerprints by mapping (1 if contact, 0 otherwise) either residue pairs at each vector position (Complex fingerprints, or CF), or contact positions separately for the receptor and G-protein (Receptor and G-protein fingerprints, respectively RF and GF). We performed unsupervised, hierarchical clustering on rows (unique GPCR-G-protein complex) using the *“Ward”* method and Euclidean distance as metric, employing the *clustermap* function from *seaborn* library (version 0.11.1). Rows are color annotated to indicate: G-protein bound in the experimental structure, GPCR class, experimental reported couplings (according to UCM). Top plot indicates the enrichment of the contacts observed at each position.

For each consensus position of GPCR and Gprotein we calculated the log-odds ratio (LOR) from the following contingency Table 1:
using the following equation (1):

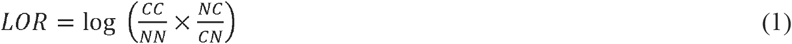

**Table 1.** Contingency table for calculating log-odds ratio

| Contact pair/G-protein | Contact | No contact |
| --- | --- | --- |
| Coupled | CC | CN |
| Not coupled | NC | NN |

*CC* and *CN* terms represent the number of GPCRs coupled to a specific Gprotein group (G_i/o_ or G_s_) that are or are not, respectively, a contact at that position (either individual GPCR or G-protein positions or residue pairs). *NC* and *NN* terms represent the number of non-coupled GPCRs for a specific G-protein, that are or are not in contact, respectively, at a given position (either individual GPCR or Gprotein positions or residue pairs). Contacts contributed from the loops, N-termini and C-termini of the GPCR where aggregated. We calculated the binning statistics of the log-odds ratio of contacts.

### Clustering of G-proteins complex conformations

We compared G_i/o_ and G_s_ complexes by performing Root Mean Square Deviation (RMSD)-based clustering. To calculate RMSD, we created a list of consensus positions based on all the sequences of GPCR and all G-protein in 264 complexes, by first mapping PDB residues to Uniprot canonical sequences via SIFTS ^39^ and then to GPCRdb consensus numbers^27^. We considered 146 GPCR and 80 Gprotein consensus positions defining respectively the consensus core of the 7TM domain and the Ras GTPase domain solved in all experimental structures. The complexes were fit using the Cα atoms of the GPCR core, while the Cα of the G-protein core where used to calculate the RMSD after superimposition. Calculations where performed using the Superimposer function of the PDB Biopython module^43^ (version 1.78) through customized scripts. We performed hierarchical clustering on RMSD using the Ward method with Euclidean distance as metrics, using the *clustermap* function from *seaborn* library (version 0.11.1). We compared the distribution of the RMSD calculated among complexes of the G_i/o_ and G_s_ groups using a Wilcoxon rank-sum test. Results were displayed through matplotlib (https://matplotlib.org/) and seaborn (https://seaborn.pydata.org/) libraries using customized python scripts. We also calculated the root mean squared fluctuations of the G-protein consensus positions using the following equation:

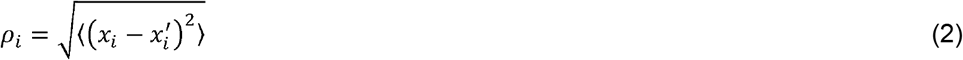

where *x_i_* is the coordinate of particle *i* and 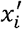 is the coordinate of particle *i* in the reference structure ‘, which is the complex with the least RMSD deviation from the other complexes (i.e. centroid) in the G_s_ (PDB: 6p9x) and _Gi/o_ (PDB: 7y15) groups.

We compared G_s_ and G_i/o_ groups RMSFs by performing a Wilcoxon test and we plotted each position and its standard deviation.

### Analysis of GPCR-Gα subunit binding energy with Rosetta

To analyze the GPCR-Gα interface in a 3D structural model, we first relax the structure using the Rosetta relax application ^44^, using backbone constraints, to eliminate small clashes that sometimes can be found in experimental structures and that can affect the Rosetta energy function. Then we run Rosetta InterfaceAnalyzer^31^, from RosettaCommon software suite (version 2021.16.61629), specifying the chains of the GPCR and the Gα which are interacting in the complex. This protocol takes a multichain complex as input and computes a new structure in which the two chains of interest are separated. The interface energy and the ΔSASA are calculated as the difference in energy and SASA in the bound and unbound structure. We run InterfaceAnalyzer with the -pack_input and -pack_separated flags to optimize the side chain configuration before and after separating the chains. If a nanobody was present in a structure, we removed it before the relaxation step, to limit its influence on the analysis.

The interface energy is computed according to the Rosetta energy function, which includes physics-based terms that represent electrostatic and van der Waals’ interactions, as well as statistical terms representing the probability of finding the torsion angles in the Ramachandran plots. This score is indicated in Rosetta Energy Units (REU) and cannot be converted into the actual binding energy, but it gives a reasonable estimation of the stability of the complex^45^.

## Availability

Code and data used for this study are available at: *https://github.com/raimondilab/GPCR_structure_analysis*

## Acknowledgments

We gratefully acknowledge the CINECA award, in collaboration with AIRC, for the availability of high performance computing resources and support. We gratefully acknowledge computational resources of the Center for High Performance Computing (CHPC) at Scuola Normale Superiore.

## Supplementary Figures

**Figure S1:** secondary couplings from UCM of Gs (left) and Gi/o coupled receptors. Bars indicate the value as a fraction over the total number of receptors in the UCM dataset.

**Figure S2:** GPCR-G-protein contact interface fingerprint (or CF fingerprint): each row is a GPCR-G-protein contact positions (referenced respectively to GPCRdb numbering and G-protein position (CGN) numbering) and each column is a unique receptor. If a receptor is complexed with more than one G-protein, its complex fingerprint is reported accordingly. Column are color annotated to indicate: G-protein bound in the experimental structure, GPCR class, experimental reported coupling (according to UCM, or either GEMTA, Shedding or GtoPDB). Only contacts present in at least 20% of the structures are shown, considering all unique complexes with the four G-protein families.

**Figure S3:** A) receptor and B) G-protein fingerprint clustering. Each column is a GPCR (or G-protein) contact positions (referenced respectively to GPCRdb and G-protein position (CGN) numbering) and each row is a unique receptor. If a receptor is complexed with more than one G-protein, its complex fingerprint is reported accordingly. Rows are color annotated as Fig. S2’s columns

**Figure S4:** RMSD hierarchical clustering for all the 264 structures analysed. Color annotations are as in Figure 4A

**Figure S5:** A) RMSD hierarchical clustering of representative structures of unique complexes of Class A receptors. B) RMSD distributions of G_s_ and G_i/o_ complexes; statistics is performed via a Wilcoxon rank-sum test

**Figure S6:** A) RMSD hierarchical clustering of representative structures of unique complexes after Rosetta relaxation. B) RMSD distributions of G_s_ and G_i/o_ complexes; statistics is performed via a Wilcoxon rank-sum test

**Figure S7:** A) RMSD hierarchical clustering of representative structures of unique complexes of Class A receptors after Rosetta relaxation. B) RMSD distributions of G_s_ and G_i/o_ complexes; statistics is performed via a Wilcoxon rank-sum test

**Figure S8:** binding energy estimated through Rosetta for class A receptor bound to G_s_ and G_i/o_ proteins A) ΔG binding (RU); B) Delta Solvent Accessbile Surface Area (ΔSASA); C) ΔSASA/ΔAG.

## Supplementary Tables

**Table S1:** list of PDB structures considered for the study

**Table S2:** chains corresponding to G protein from representative PDB and residue number mapping to Common G protein numbering

**Table S3:** chains corresponding to GPCRs from representative PDB and residue number mapping to GPCRdb numbering

**Table S4:** binary GPCR-Gα subunit complexes predicted through AlphaFold-multimer with Rosetta binding energy (InterfaceAnalyzer) and structural filter annotations

